# Environmental Conditions and Vigilance Behaviour in the Smooth-coated Otter, *Lutrogale perspicillata*

**DOI:** 10.1101/2022.03.21.485127

**Authors:** N. P. Travell, K. Fernandez

## Abstract

Vigilance is an ecologically important behaviour that acts as a defence mechanism against threats such as predators, competitors, and environmental pressures. Being vigilant can place energetic constraints on an individual, as conducting other behaviours such as grooming can be limited. The smooth-coated otter is a vulnerable species, and understanding baseline behaviour, such as vigilance, and stressor effects can be a crucial step in implementing any conservation strategies. However, literature on this charismatic species is relatively sparse, with the majority of research covering prey items, habitat use and extent of occurrence.

Using camera trap footage along with historical climate datasets, the current study assessed various aspects of vigilance behaviour of the smooth-coated otter. These aspects include how long individuals dedicated to being vigilant, what they prioritised being vigilant over, and how intense their vigilance efforts were

Variables that are expected to change with global climate change such as temperature, and to an extent, air pressure, were associated with increases in vigilance duration. Wind speeds, which are also expected to be affected by climate change, along with increasing numbers of conspecifics, were linked to increased intensity possibly highlighting that increasing ambient movement results in more frequent head rotations. When vigilance is ineffectual however, such during periods of higher rainfall, cloud cover or at night, intensity decreases, and gazing towards conspecifics increases - possibly to reaffirm where safety lies when threats cannot be adequately monitored.

## 1 Introduction

The smooth-coated otter (*Lutrogale perspicillata*) is one of four species of otter found in Asia and is typically found in groups of 1-9 (Hussain 1997, Bungum et al 2021) within the waterways of Asia (Estes 1986). They are classified as vulnerable by the IUCN with a decreasing population trend (de Silva et al 2015), largely due to anthropogenic influences such as habitat change (for example the production of dams) or illegal collection occuring due to the pet trade or for their fur (Gomez et al 2016). As apex predators, they improve biodiversity through the control of prey species populations (Ritchie and Johnson 2009). This includes the predation of invasive species (Wallach et al 2015) which, when considering the geometrically increasing rate of introduced species occurrence (Seebens et al 2017), highlights the need to protect the smooth-coated otter.

Prior to the implementation of any protection strategies, a thorough understanding of baseline physiology and behaviours, along with an appraisal of stressors and their effects, is required (Gessner and Rochard 2011, Stevenson 2006). Though otters are considered charismatic ‘flagship’ fauna (Stevens et al 2011), literature on baseline behaviours remains relatively scarce, with the majority of recent literature covering the status of the species and extent of occurrence, and less commonly habitat use or diet. One such behaviour of high ecological importance that acts as a defence mechanism for predation and threat assessment, is vigilance.

### 1.2 What is vigilance behaviour?

Animals use various senses to monitor their surroundings for potential threats (Galton 1871). Upon detection alarm calls can be made, alerting conspecifics to the threat, and allowing them time to adopt anti-predator behaviours such as defensive formations (Beauchamp 2015). Vigilance behaviour occurs within apex predators as well, despite not being directly predated upon, as intra-guild competition can pose a threat to their fitness if other individuals are better able to exploit their resources (Caro and Stoner 2003, Eaton 1979, Kimbrell et al 2007). Additionally, lower ranked individuals within a group can be more vigilant around threatening conspecifics (Kraus et al 2011), and extreme environmental conditions such as cyclones and flash floods that can present both threats and opportunities. Furthermore, the increased human-otter conflict arising from competition for the same resources, ie territory and food, or the exploitation of wild specimens for the pet/pelt trade also drives an increase in vigilance (Ciuti et al 2012).

### 1.3 Why vigilance?

Vigilance has long been considered a costly activity (Lima & Dill, 1990), as time spent being vigilant detracts from time spent on other fitness-increasing, or sustaining, activities such as foraging (though see Fortin et al 2004 and Cowlishaw et al 2004). The cost of vigilance is not held by the vigilant individual alone however, as thwarting a threat such as predation will incur a cost in the predator. Conversely vigilance may not be costly to the individual at all (Illius and Fitzgibbon 1994), or may depend on whether the purpose of the behaviour is for predator avoidance or social monitoring (Alberts 1994, Gaynor and Cords 2012). Assessment of gaze direction can help clarify what the vigilant individual sees as a threat. Additionally, vigilance may be difficult to distinguish - for instance an individual may have their head up without being intentionally vigilant - so activity such as head rotations, which incur a mechanical cost, can also be assessed to highlight the intensity of which an individual is being vigilant.

In addition to vigilance being thought of as a state of behaviour, it can also be considered a state of information processing with inherent qualitative costs. For instance, vigilance can have subtle effects within the eyes, such as the widening of eyelids (Sussking et al 2008), lowered blinking rate (Matsumoto et al 2018, Cross et al 2013, Yorzinski 2016), and pupil dilation (Ebitz et al 2014). These serve to increase the rate of information intake, such that the crucial milliseconds during which a threat appears won’t be missed, however this results in a reduction of visual acuity (Ebitz et al 2014). Additionally, this high rate of information processing is difficult to sustain (Dukas and Clark 1995, McIntire et al 2014), and the resulting vigilance decrement can lead to individuals being unable to detect predators or obscured food items. Therefore, vigilance, and the rate of vigilance recovery, can act as a dominant factor regarding animal decision making (Dukas and Clark 2014).

Using camera trap footage along with historical climate datasets, the current study assessed various vigilance behaviour aspects of the smooth-coated otter, such as vigilance duration, intensity, and purpose, by constructing time budgets, assessing head rotations, and gaze direction.

## 2 Methods

### 2.1 Data collection and study location

Remote camera traps with passive infrared sensors were set up at two different locations in Tiswadi Taluka of the state of Goa in South India. Each location had a different habitat used by the smooth-coated otter. A single camera was set up at a defecation site frequented by a family of otters located on the Island of Chorao during February and March 2021. This site is located along the River Mandovi. The second site, a natal den site frequented by a different family, is located in the Salvadore Wetlands along the River Mapusa. A single camera was placed here over the course of the study but at different angles in the month of November in 2018 and 2019. The camera traps were set to take 30 second videos per trigger with a one second gap between consecutive triggers.

### 2.2 Vigilance activity budget collection method

To begin with, all observed otter behaviour from the data set have been defined and presented in an ethogram (Table 1). For the purposes of this study, a specific data set was chosen based on ‘suitable clips’ which is defined as an observation of an individual otter on screen, that exhibited vigilance behaviour as defined in the ethogram (Table 1). An activity budget was created to include only vigilance and was calculated as a percentage of time that the otter was within the footage. Only clips where the face of the otter was fully visible were included, or when whiskers could be used to infer head position with confidence. Seconds of vigilance duration were only counted when the head was not faced downwards, such as during foraging behaviour, and the animal appeared to be outwardly alert. In total 303 clips were deemed suitable, for a total of 4555 seconds of otter behaviour.

| Behaviour | Description |
| --- | --- |
| Resting | Individual is lying down relatively motionless. Their eyes can be open or closed. |
|  | but their head is not up in a vigilant posture, and they are not looking around. |
| Locomotion | Individual is crawling, swimming (with either the head up or diving underwater), or jumping |
| Foraging | Movement with no direct route used, and head is mostly down, nose into the ground, possibly with prey item in sight. Paws can be used to move vegetation, dig, or claw the ground. |
| Socialising | Individual is interacting with another, this can consist of carrying the offspring in the adult's jaws, grooming, vocalising, nudging, and play fighting. |
| Excretion | Individual is engaged in excretion of either urine or faecal matter. Typically the head and tail are raised, and the body is in a convex, standing, manner. |
| Grooming | The individual is using their mouth or paws to clean themselves or scraping their skin against a surface. Often includes rolling around |
| Nest maintenance | Individual is carrying nesting material back and forth from the den |
| Vigilance | Being alert, looking around while stationary. The head and ears can point towards a focal point of interest |

### 2.3 Vigilance intensity collection method

Head rotation data collection happened simultaneously to the vigilance activity budget and included monitoring how many times a change in head position occurred during exhibition of vigilance behaviour. Head rotations were recorded only if a distinct positional change (>=15 degrees) occurred with a stop - slowly moving the head was not accounted for unless the head was fixed on a position. Rotations were not counted if the individual was changing position during another behaviour such as for grooming behaviour.

### 2.4 Vigilance focal points collection method

To further elucidate the purpose of vigilance under different environmental conditions, the focal points of each otter was also collected. Due to the difficulty of assessing accurate focal locations from a 2D recording, general gaze direction was assessed, including recurring and easily recognisable landmarks such as the waterline, sky, inland and conspecifics. Parallel to the intensity collection method, focal points were only recorded when a distinct head position occurred. Areas that the otter looked at were tallied up, then a percentage budget was constructed.

### 2.5 Environmental Datasets

To determine how otter vigilance behaviour varies with the environment, climatic datasets were extracted (https://www.worldweatheronline.com/). Factors such as temperature (degrees celsius °C), cloud cover (%), air pressure (millibars mb), visibility (kilometres km), and rainfall (mm) were included. However due to the ambiguity of recording date:time, averages were taken over the duration that the camera was in operation. Additionally, wind speed was subtracted from gust speed to create a gust disparity factor (kilometres per hour), highlighting which days had the largest differences between sudden high gust speeds and ambient wind flow.

### 2.6 Statistics

For statistical analysis, R Studio (Version 1.4.1717) was used, along with the following packages: diplyr, tidyr, gridExtra, and ggplot2. Statistical tests were run with and without outliers, and the significance of the tests did not change. Results presented in this paper are listed with outliers in the dataset, unless otherwise stated.

## 3 Results

### 3.1 Vigilance activity budget

Otters were vigilant 36.09% percent of the time (SD+/-34.99), which differed by site (F(1,286)=31.5, p=4.71e-08), month (F(2,285)= 15.81, p=<0.001), year (F(2,285)=22.19, p=<0.001), and whether the clip was recorded at night or day time (F(1,286)= 24.7, p=<0.001). Additionally, certain behaviours require the individual to move their head, E.G. away from the vigilant position and towards the ground as seen during foraging behaviour, in this case vigilance duration was seen to differ significantly per concurrent behaviour (F(7,280) =-6.67, p=<0.001) (see figure 1).

**Figure 1.**
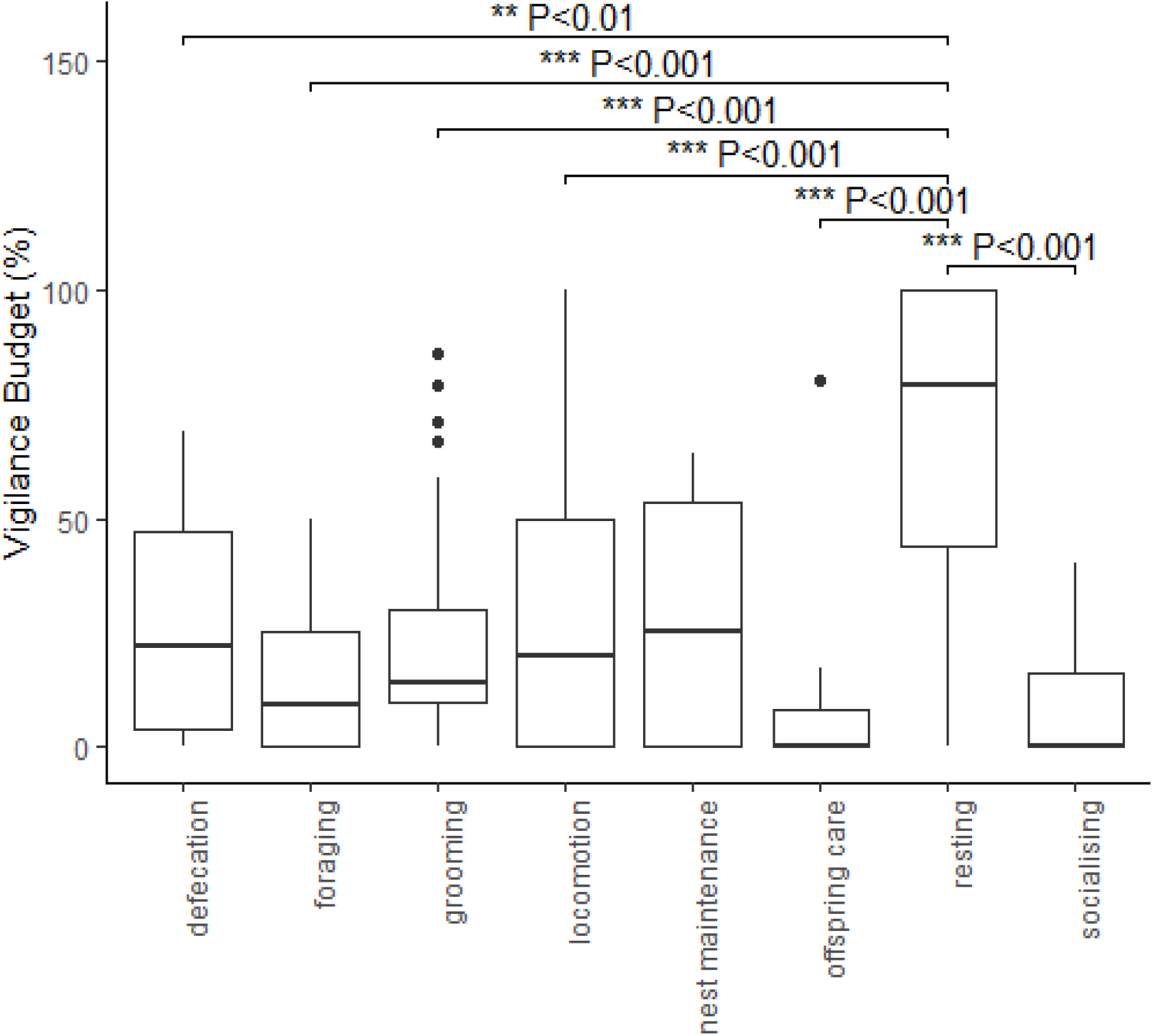
Boxplots showing the distribution of vigilance duration (%) and how it changes by concurrent behaviour. Significance values were determined by TukeyHSD post-hoc comparisons

### 3.2 Vigilance intensity

Head rotation (calculated as the frequency of head rotations that occurred during the total time the individual was seen to be exhibiting vigilance behaviour) data was found to be non-normal, so was transformed using the following formula: log(rotations/s +1). The transformed variable was normally distributed. The following statistically significant relationships with environmental factors were found: Site (F(1,286)=31.5, p= <0.001), main behaviour seen by the individual during the clip (F(7,280)-6.67,p=<0.001), whether the clip was recorded at night or day time (F(1,286)=24.7,p=<0.001), month of recording (F(2,285)=15.81,p=<0.001), and year (F(2,285)=22.19,p=<0.001).

### 3.3 Vigilance focal points

Smooth-coated otters were seen to pay unequal attention to their surroundings, with a difference occurring in focal gaze location (F(5,936)=68.98,p=<0.001).A square root transformation was applied to the percentage of time the individual spent gazing at each direction in order to achieve a normal distribution, and the results of focal direction analysed using a parametric MANOVA. Site type (F(1,5)=2.557, p=0.110), time of day (F(1,5)=0.468, p=0.494), year (F(2,5)=1.281, p=0.278), month (F(2,5)=1.711,p=0.181), and behaviour (F(7,5)=3.6, p=0.789) alone was not seen to significantly change focal direction, however an interaction effect was noted for all of these variables - site x direction (F(1,5)=14.307, p=<0.001), time of day x direction (F(5,1)=3.889,p=0.002), year x direction (F(2,10)=7.572,p=<0.001), month x direction (F(2,10)=7.565,p=<0.001), and behaviour x direction (F(7,35)=21.2,p=<0.001). Subsequent post-hoc comparison tests (TukeyHSD) did not reach significance.

## 4 Discussion

Though individual variables had poor explainability (as expressed as adjusted R^2^) over variation in vigilance behaviour (with the highest being 13.8% for both visibility on head rotations per second, and wind speed on vigilance budget), statistical significance was found for many relationships when assessing the association between environmental variables and the three measured aspects of vigilance behaviour.

### 4.2 Vigilance budget

A positive trend for vigilance duration and the numerical variables was only seen for temperature and air pressure (table 3). Increases in temperature is expected as part of global climate change, and sea level air pressure was associated with global warming in the Indian Ocean region (Copsey et al 2006), possibly resulting in future drivers of smooth-coated otter vigilance behaviour. Changes in air pressure has long been known to be associated with a variety of behavioural changes in a range of animal species, such as increasing pain sensitivity in rats (*Funakubo et al 2010*), calving incidence in domestic cows (*Dvorak 1978*), and birds engaging in faster foraging periods (Metcalfe et al 2013), so the change in vigilant behaviour noted within the current study is not totally unexpected.

**Table 2.** depicting how the mean vigilance budget (%) varies by level of categorical factor

| Predictor Category | Predictor | Mean |
| --- | --- | --- |
| Site type | Defecation | 21.3±22 |
|  | Den | 44±38 |
| Time of day | Day | 35.0±32.9 |
|  | Night | 37.9±38.3 |
| Year | 2018 | 50.3±38.1 |
|  | 2019 | 32.7±35.3 |
|  | 2021 | 21.3±22 |
| Month | February | 19±23.4 |
|  | March | 22.2±21.5 |
|  | November | 44±38 |

**Table 3.** depicting linear statistics of the relationships between vigilance duration and the continuous variables

| Variable | Adjusted R2 | Intercept | Slope | Significance |
| --- | --- | --- | --- | --- |
| Conspecifics Present | 0.0241 | 47.421 | -10.23 | <0.001 |
| Air Pressure | 0.09277 | -12407.17 | 12.309 | <0.001 |
| Rainfall | 0.02327 | 118.7248 | -0.4237 | 0.005 |
| Wind Speed | 0.1384 | 110.712 | -7.515 | <0.001 |
| Gust Speed | 0.1352 | 103.3886 | -4.201 | <0.001 |
| Gust Disparity | 0.1226 | 90.45 | -8.926 | <0.001 |
| Cloud Cover | -0.003111 | 38.0303 | -0.1272 | 0.741 |
| Visibility | 0.02024 | 735.51 | -70.57 | 0.009 |
| Temperature | 0.1108 | -476.431 | 17.501 | <0.001 |

**Table 4.**
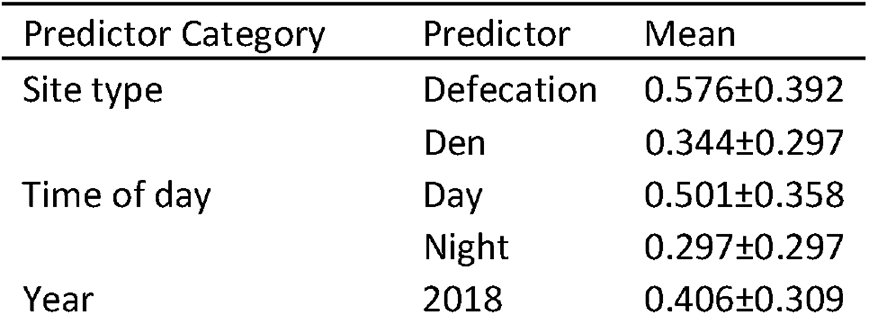
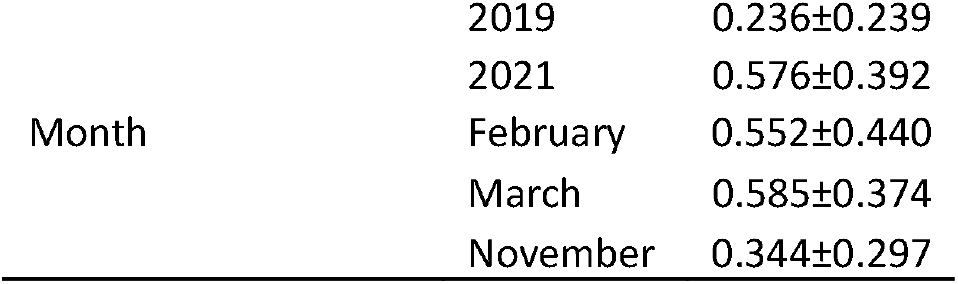
depicting how the mean log+1 transformed rotations per second varies by level of categorical factor

**Table 5.**
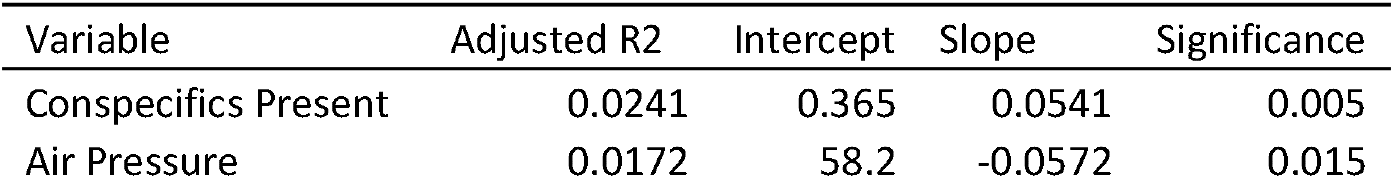
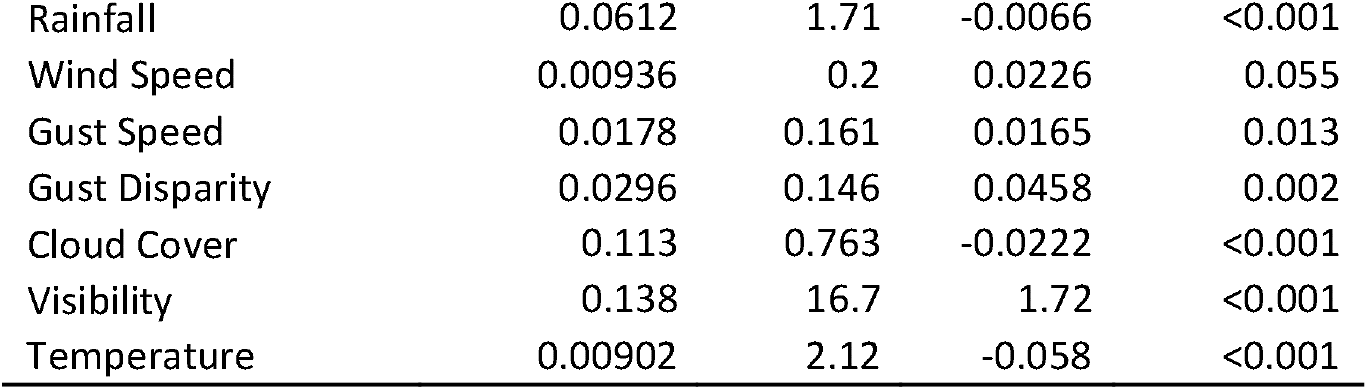
depicting linear statistics of the relationships between transformed head rotations per second and the continuous variables

**Table 6.** depicting how the mean time budget varies by focal direction

| Direction | Mean |
| --- | --- |
| At a Conspecific | 6.94±12.9 |
| Down | 17.3±19.9 |
| Inland | 38.7±27.3 |
| Sky | 6.46±14.3 |
| Waterline | 6.16±13.5 |
| Unknown | 22.5±23.9 |

**Table 7.**
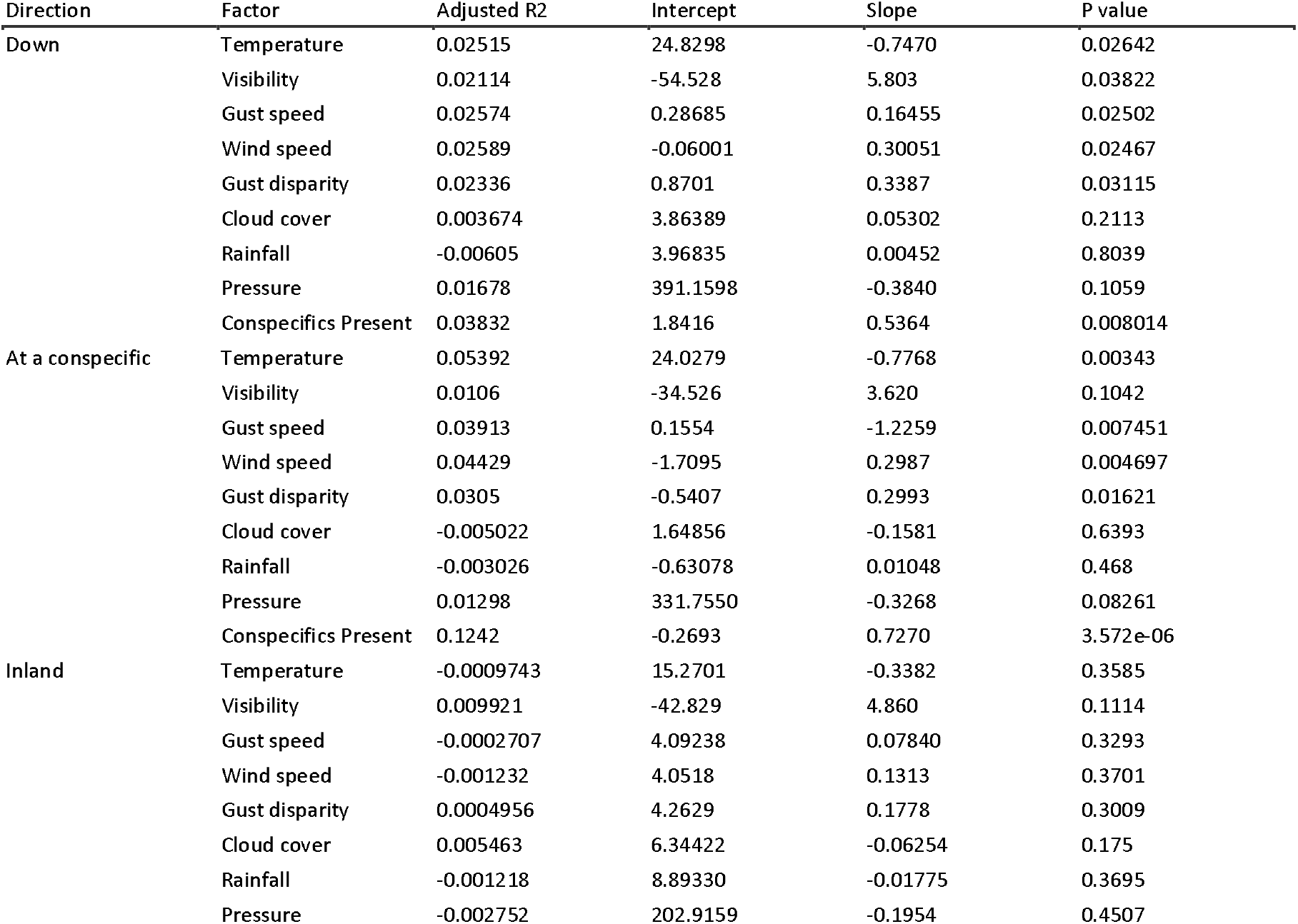
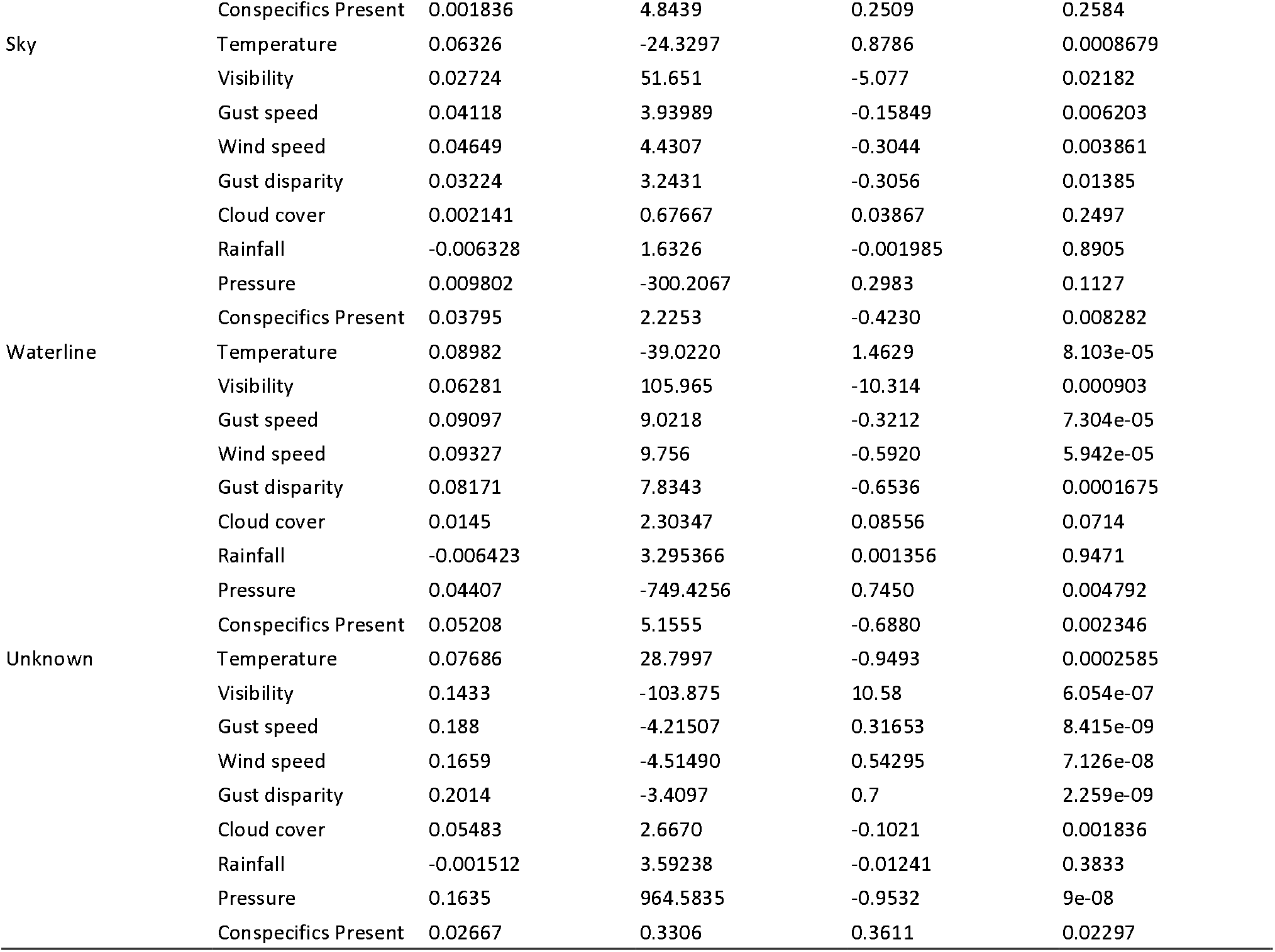
depicting linear statistics of the relationships between the percentage of time otters spent looking in each direction and the continuous variables

Concurrent behaviours may affect vigilance budget more than the other measured aspects of vigilance behaviour due to the inherent requirement of the body to be used in a certain manner during some of these behaviours. For example, the head would need to be in an upwards ‘vigilant’ position during nest maintenance else the nesting material would drag on the ground, and the head would need to be facing downward in a non-vigilant manner during foraging behaviour. As visualised in figure 1, the concurrent behaviours and their effects could be grouped into two categories: those associated with increased vigilance - resting (mean 66.9% vigilance duration), locomotion (28.9%), nest maintenance (28.5%), and defecation (27.1%) - and those associated with lower vigilance budgets - grooming (22.2%), foraging (14.5%), offspring care (12%), socialising (8.62%).

### 4.3 Vigilance intensity

Head rotations per second were associated with ambient movement around the vigilant individual, such as that caused by increasing numbers of conspecifics present or with increased wind speeds (which are also expected to rise due to global climate change). The slope for gust disparity (the difference between gust speed and wind speed) was steeper than both wind and gust speed, indicating that more sudden movements result in more frequent gaze direction changes. When vision was restricted however, such as during periods of higher rainfall or cloud cover, or at night, then lower rotations per second were noted. A similar relationship was present between rainfall and vigilance duration - possibly highlighting that in scenarios where vigilance would be ineffectual, an individual may adaptively reduce vigilance and choose to spend their time on other activities

Though not directly competitive with other activities like vigilance duration, head rotations per second are still influenced by the position of the body and thus are affected by the concurrent behaviour seen. However, there seems to be less variation between the behaviours, with only grooming being significantly different. When grooming, an individual will have their head in a downwards position, making ‘accidental’ vigilance impossible and requiring faster head rotations in order to adequately scan the environment (as seen in figure 2).

**Figure 2.**
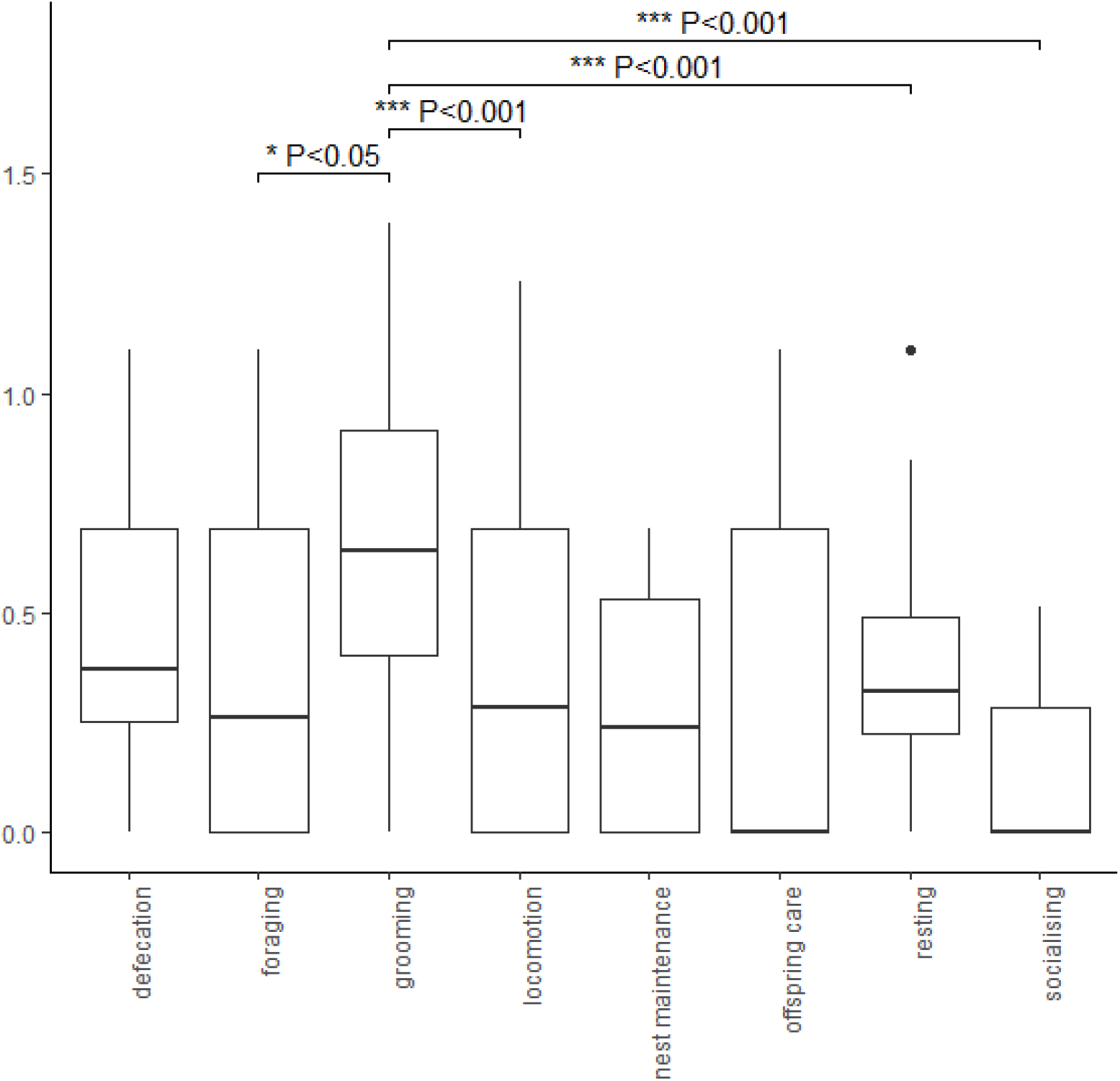
depicting the variation of head rotations per second (once transformed to achieve normal distribution) by concurrent behaviour

**Figure 3-11:**
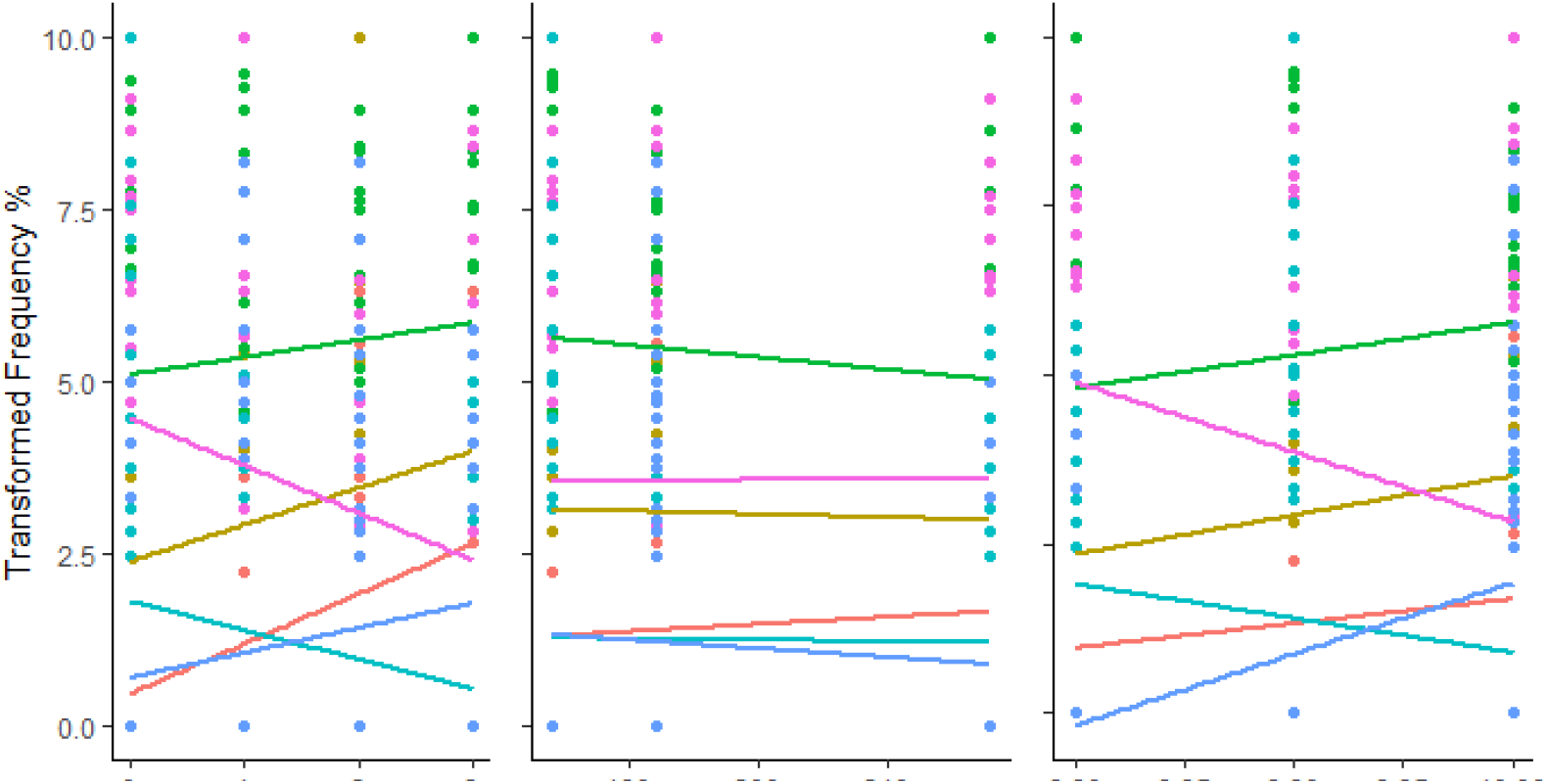
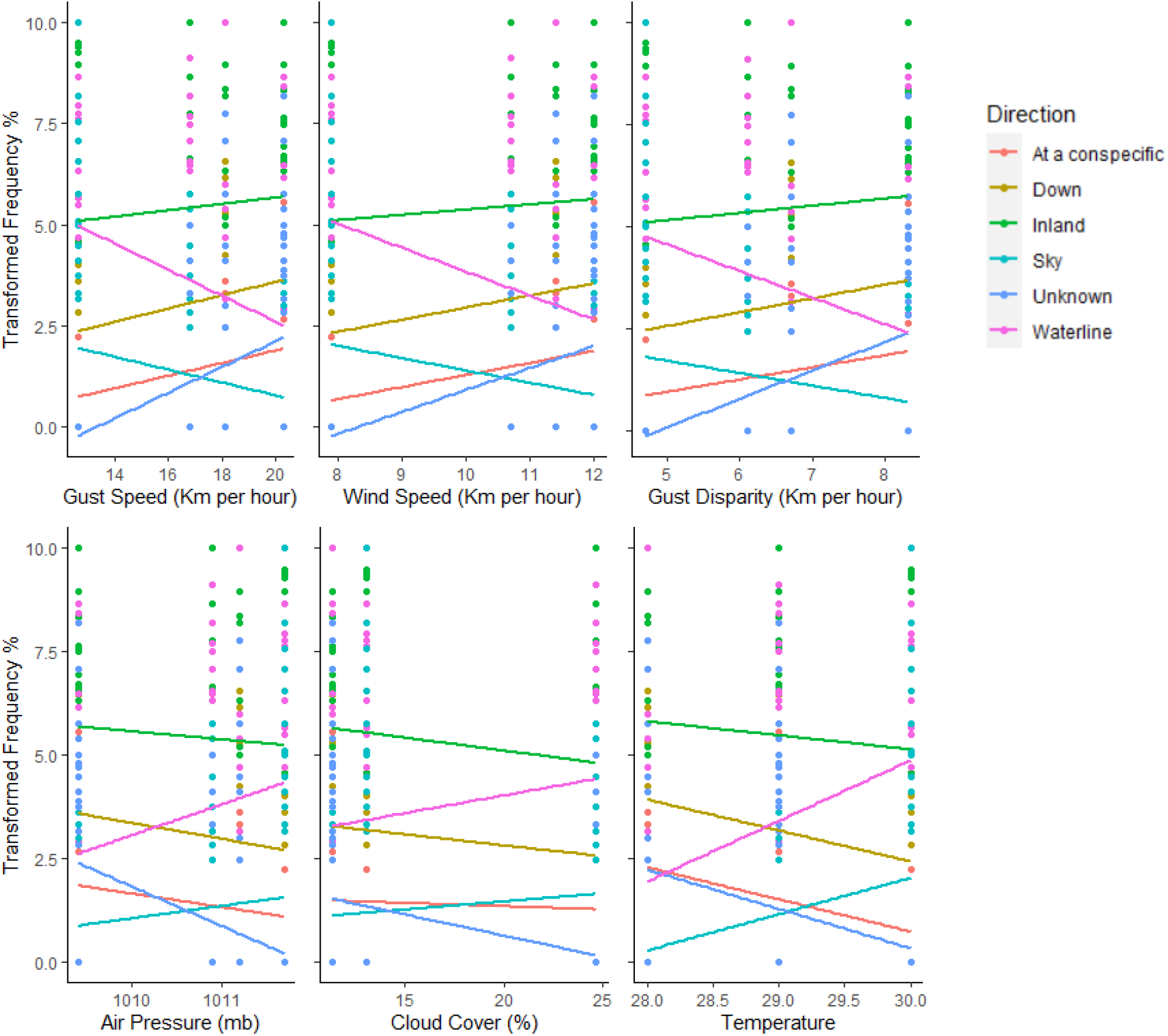
Scatter plots showing the relationships between the percent of time otters spent gazing in each direction (colours) and environmental variables (figures). A square root +1 transformation was applied to the frequency budget to achieve normality. See table 7 in appendix for regression equations and significance.

### 4.4 Vigilance focal points

As wind speeds and head rotations per second were increasing, the smooth-coated otters were increasingly gazing inland. Noting the earlier relationship between ambient movement and head rotations, a similar causal relationship may be present here; floral matter would be influenced by the wind and thus create sudden movements to be inspected. In contrast to this, increasing wind and gust speeds were associated with a lower frequency of waterline gazing - possibly due to the disruption on the water surface making any vigilance attempts less productive. Furthermore, when vigilance is made ineffectual due to rainfall, otter gazing was directed more towards conspecifics, which may be a strategy to reaffirm where relative safety lies when threats are less effectively observed.

### 4.5 Further research

Several climatic factors expected to be affected by global climate change were associated with increases in vigilance duration, vigilance intensity, and variation in gaze direction. Though behavioural and physiological flexibility of the smooth-coated otter may allow adaptation (Arenz and Leger 2000, Killen et al 2015, Uchda et al 2019), it is possible this increased demand on an individual’s energetic constraints could result in a poorer body condition (due to less foraging, grooming, and other activities) and thus reducing individual fitness (Lima & Dill, 1990). Further research should be conducted to determine the limits of otter vigilance, and to clarify what effects variation in vigilance behaviour has on body condition.

Due to memory card issues, the exact time and date of each clip was unreliable, and thus correlating the behaviour seen to precise climate records was not possible. Furthermore, due to the inherent weakness of a single camera trap, identifying where the otter was looking beyond a rough direction from a 2D plane was not possible. Follow up field surveys rectifying this will allow for more reliable inferences to be drawn between climatic factors and vigilance behaviour, along with highlighting what the smooth-coated otters prioritise being vigilant over.

## 6 Conclusion

In the current study, the relationship between climatic variables and smooth-coated otter vigilance behaviour was assessed using camera trap footage. The results in this paper can be grouped into three hypotheses regarding smooth-coated otter vigilance. Whether vigilance will be ineffectual (such as through rainfall reducing visibility, or wind speeds obscuring water surfaces) seemed to be associated with where otters directed their gaze, whilst ambient movement (particularly sudden spikes occurring in part due to neighbouring conspecifics or gusts) may drive head rotations per second. Finally, climate change may be a driver of increased vigilance duration, through increased temperatures (and possibly air pressure). Follow up studies were recommended to both clarify the link between vigilance and body condition within the species, and confirm the accuracy of the noted trends.

## 7 Acknowledgements

This research did not receive any specific grant from funding agencies in the public, commercial, or not-for-profit sectors.

## 8 Conflict of Interests

The authors declare no competing interests.

## 7) Appendix

## Notes

### Competing Interest Statement

The authors have declared no competing interest.

## References

Acolas M.L., Gessner J., & Rochard E. (2011). Population Conservation Requires Improved Understanding of In Situ Life Histories. In: Williot P., Rochard E., Desse-Berset N., Kirschbaum F., Gessner J. (eds) Biology and Conservation of the European SturgeonAcipenser sturioL. 1758. Springer, Berlin, Heidelberg. https://doi.org/10.1007/978-3-642-20611-5_44

Alberts, S. (1994). Vigilance in young baboons: effects of habitat, age, sex, and maternal rank on glance rate. Animal Behaviour, 47, 749e755.

Arenz, C.L. and Leger, D.W. (2000). Antipredator vigilance of juvenile and adult thirteen-lined ground squirrels and the role of nutritional need. Animal Behaviour, [online] 59(3), 535–541.

Beauchamp, G. (2015). Animal vigilance : monitoring predators and competitors (1st ed.). Boston: Academic Press.

Bungum, H., Yee, M., Borker, A., Hsu, C., & Johns, P. (2021). Multiple reproductive females in family groups of smooth-coated otters. BioRxiv doi: 10.1101/2021.06.26.450053

Caro, T., & Stoner, C. (2003). The potential for interspecific competition among African carnivores. Biological Conservation, 110(1), 67–75. doi: 10.1016/s0006-3207(02)00177-5

Ciuti, S., Northrup, J., Muhly, T., Simi, S., Musiani, M., Pitt, J., & Boyce, M. (2012). Effects of Humans on Behaviour of Wildlife Exceed Those of Natural Predators in a Landscape of Fear. Plos ONE, 7(11), e50611. doi: 10.1371/journal.pone.0050611

Copsey, D., Sutton, R., & Knight, J. (2006). Recent trends in sea level pressure in the Indian Ocean region. Geophysical Research Letters, 33(19), L19712, https://doi.org/10.1029/2006GL027175

Cowlishaw, G., Lawes, M., Lightbody, M., Martin, A., Pettifor, R., & Rowcliffe, J. (2004). A simple rule for the costs of vigilance: empirical evidence from a social forager. Proceedings Of The Royal Society Of London. Series B: Biological Sciences, 271(1534), 27–33. doi: 10.1098/rspb.2003.2522

Cross, D., Marzluff, J., Palmquist, I., Minoshima, S., Shimizu, T., & Miyaoka, R. (2013). Distinct neural circuits underlie assessment of a diversity of natural dangers by American crows. Proceedings Of The Royal Society B: Biological Sciences, 280(1765), 20131046. doi: 10.1098/rspb.2013.1046

de Silva, P., Khan, W.A., Kanchanasaka, B., Reza Lubis, I., Feeroz, M.M. & Al-Sheikhly, O.F. (2015). Lutrogale perspicillata. The IUCN Red List of Threatened Species 2015: e.T12427A21934884. https://dx.doi.org/10.2305/IUCN.UK.2015-2.RLTS.T12427A21934884.en.

Dukas, R., & Clark, C. (1995). Sustained vigilance and animal performance. Animal Behaviour, 49(5), 1259–1267. doi: 10.1006/anbe.1995.0158

Dvorak, R. (1978). A note on the relationship between barometric pressure and calving incidence. Animal Reproduction Science, 1(1), pp.3–7.

Eaton, R.L., (1979). Interference competition among carnivores: a model for the evolution of social behavior. Carnivore, 2(9–16), 82–90.

Ebitz, R., Pearson, J., & Platt, M. (2014). Pupil size and social vigilance in rhesus macaques. Frontiers In Neuroscience, 8. doi: 10.3389/fnins.2014.00100

Estes, J. (1986). Riverine Mammals: Otters. Ecology and Conservation. Science, 233(4770), 1333–1334. doi: 10.1126/science.233.4770.1333.b

Fortin, D., Boyce, M., Merrill, E., & Fryxell, J. (2004). Foraging costs of vigilance in large mammalian herbivores. Oikos, 107(1), 172–180. doi: 10.1111/j.0030-1299.2004.12976.x

Funakubo, M., Sato, J., Obata, K. and Mizumura, K., 2010. The rate and magnitude of atmospheric pressure change that aggravate pain-related behavior of nerve injured rats. International Journal of Biometeorology, 55(3), pp.319–326.

Galton, F. (1871). Gregariousness in cattle and in men, Macmillan’s Magazine, 23 (136), 353–357

Gaynor, K. & Cords, M. (2012). Antipredator and social monitoring functions of vigilance behaviour in blue monkeys. Animal Behaviour 84, 531–537, https://doi.org/10.1016/j.anbehav.2012.06.003

Gomez, L., Leupen, B., Theng, M., Fernandez, K., & Savage, M. (2016). Illegal Otter Trade : An analysis of seizures in selected Asian countries (1980-2015). TRAFFIC.

Hussain, S., & Choudhury, B. (1997). Distribution and status of the smooth-coated otter Lutra perspicillata in National Chambal Sanctuary, India. Biological Conservation, 80(2), 199–206. doi: 10.1016/s0006-3207(96)00033-x

Illius, A. W., & Fitzgibbon, C. (1994). Costs of vigilance in foraging ungulates. Animal Behaviour, 47(2), 481–484. https://doi.org/10.1006/anbe.1994.1067

Killen, S.S., Reid, D., Marras, S. and Domenici, P. (2015). The interplay between aerobic metabolism and antipredator performance: vigilance is related to recovery rate after exercise. Frontiers in Physiology, 6.

Kimbrell, T., Holt, R., & Lundberg, P. (2007). The influence of vigilance on intraguild predation. Journal Of Theoretical Biology, 249(2), 218–234. doi: 10.1016/j.jtbi.2007.07.031

Kraus, M., Horberg, E., Goetz, J., & Keltner, D. (2011). Social Class Rank, Threat Vigilance, and Hostile Reactivity. Personality And Social Psychology Bulletin, 37(10), 1376–1388. doi: 10.1177/0146167211410987

Lima, S., & Dill, L. (1990). Behavioral decisions made under the risk of predation: a review and prospectus. Canadian Journal Of Zoology, 68(4), 619–640. doi: 10.1139/z90-092

Matsumoto-Oda, A., Okamoto, K., Takahashi, K., & Ohira, H. (2018). Group size effects on inter-blink interval as an indicator of antipredator vigilance in wild baboons. Scientific Reports, 8(1). doi: 10.1038/s41598-018-28174-7

McIntire, L., McKinley, R., Goodyear, C., & McIntire, J. (2014). Detection of vigilance performance using eye blinks. Applied Ergonomics, 45(2), 354–362. doi: 10.1016/j.apergo.2013.04.020

Metcalfe, J., Schmidt, K., Bezner Kerr, W., Guglielmo, C., & MacDougall-Shackleton, S. (2013). White-throated sparrows adjust behaviour in response to manipulations of barometric pressure and temperature. Animal Behaviour, 86(6), 1285–1290. doi: 10.1016/j.anbehav.2013.09.033

Ritchie, E. & Johnson, C. (2009). Predator interactions, mesopredator release and biodiversity conservation. Ecology Letters, 12, 982–998. doi:10.1111/j.1461-0248.2009.01347.x

Seebens, H., Blackburn, T., Dyer, E., Genovesi, P., Hulme, P., Jeschke, J., … & Essl, F. (2017). No saturation in the accumulation of alien species worldwide. Nature Communications, 8, 1–9. https://doi.org/10.1038/ncomms14435

Stevens, S., Organ, J.F. and Serfass, T.L. (2011) Otters as Flagships: Social and Cultural Considerations. Proceedings of Xth International Otter Colloquium, IUCN Otter Spec. Group Bull. 28A: 150–161

Stevenson, R. (2006) Ecophysiology and conservation: The contribution of energetics—introduction to the symposium. Integrative and Comparative Biology, 46(6), 1088–1092, https://doi.org/10.1093/icb/icl053

Susskind, J., Lee, D., Cusi, A., Feiman, R., Grabski, W., & Anderson, A. (2008). Expressing fear enhances sensory acquisition. Nature Neurosciences, 11, 843–850.

Uchida, K., Suzuki, K.K., Shimamoto, T., Yanagawa, H. and Koizumi, I. (2019). Decreased vigilance or habituation to humans? Mechanisms on increased boldness in urban animals. Behavioral Ecology, 30(6), 1583–1590

Wallach, A., Ripple, W., & Carroll, S. (2015). Novel trophic cascades: apex predators enable coexistence, Trends in Ecology & Evolution, 30(3), 146–154

Yorzinski, J. (2016). Eye blinking in an avian species is associated with gaze shifts. Scientific Reports, 6(1). doi: 10.1038/srep32471

